# Effect of eye globe and optic nerve morphologies on gaze-induced optic nerve head deformations

**DOI:** 10.1101/2023.12.19.572308

**Authors:** Tingting Liu, Pham Tan Hung, Xiaofei Wang, Michaël J. A. Girard

## Abstract

**Purpose:** To investigate the effect of globe and optic nerve (ON) morphologies and tissue stiffnesses on gaze-induced optic nerve head deformations using parametric finite element modeling and a Design of Experiment (DOE) approach.

**Methods:** A custom software was developed to generate finite element models of the eye using 10 morphological parameters: dural radius, scleral, choroidal, retinal, pial, peripapillary border tissue thicknesses, prelaminar tissue depth, lamina cribrosa (LC) depth, ON radius, and ON tortuosity. A 10-factor 2-level full-factorial analysis (1,024 models) was used to predict the effects of each morphological factor and their interactions on LC strains induced by 13° adduction. Subsequently, a further DOE analysis (1,024 models) was conducted to study the effects and potential interactions between the top 5 morphological parameters identified from the initial DOE study and 5 critical tissue stiffnesses.

**Results:** In the DOE analysis of 10 morphological parameters, the five most significant factors were ON tortuosity, dural radius, ON radius, scleral thickness and LC depth. Further DOE analysis incorporating biomechanical parameters highlighted the importance of dural and LC stiffness. A larger dural radius and stiffer dura increased LC strains but the other main factors had the opposite effects. Notably, a significant interaction was found between dural radius and dural stiffness.

**Conclusions:** This study highlights the significant impact of morphological factors on LC deformations during eye movements, with key morphological effects being more pronounced than tissue stiffnesses.

## INTRODUCTION

Glaucoma is one of the most common causes of blindness worldwide.^1^ The biomechanical theory of glaucoma suggests that the deformations of the optic nerve (ON) head (ONH) tissues, especially the lamina cribrosa (LC), may lead to the apoptosis of retinal ganglion cells and visual field defects, either directly or indirectly.^2^ Intraocular pressure (IOP) and cerebrospinal fluid pressure (CSFP) are the two main mechanical loads acting on the ONH that have been shown to be correlated to glaucoma pathogenesis^3–5^. Recent studies using finite element (FE) modeling,^6–8^ optical coherence tomography ^9–13^ and magnetic resonance imaging (MRI)^14–16^ have highlighted that ON traction during eye movements can yield large ONH deformations, which may be as large as or significantly larger than those caused by a substantial IOP elevation to 40 or 50 mmHg.

In vivo studies have shown that gaze-induced ONH deformations vary widely across individuals^17,18^. The differences are likely due to variations in the biomechanical properties and morphologies of the eye globe, ON and ONH. For instance, ON tortuosity varies across individuals, as shown in **Figure 1**. A less tortuous ON has been hypothesized to generate a larger traction force, and thus potentially larger ONH deformations.^15^ Therefore, to identify those who are vulnerable to ON traction during eye movements, it would be critical to identify the biomechanical and morphological factors (and their interactions) that significantly affect ONH deformations. Using FE modeling, we have previously investigated the effects of biomechanical properties on gaze-induced ONH deformations and predicted that a stiffer dura would generate a larger ONH deformation.^6^ However, the effects of eye-globe and ON morphologies on gaze-induced ONH deformations and their potential interactions with biomechanical properties remain unexplored.

**Figure 1.**
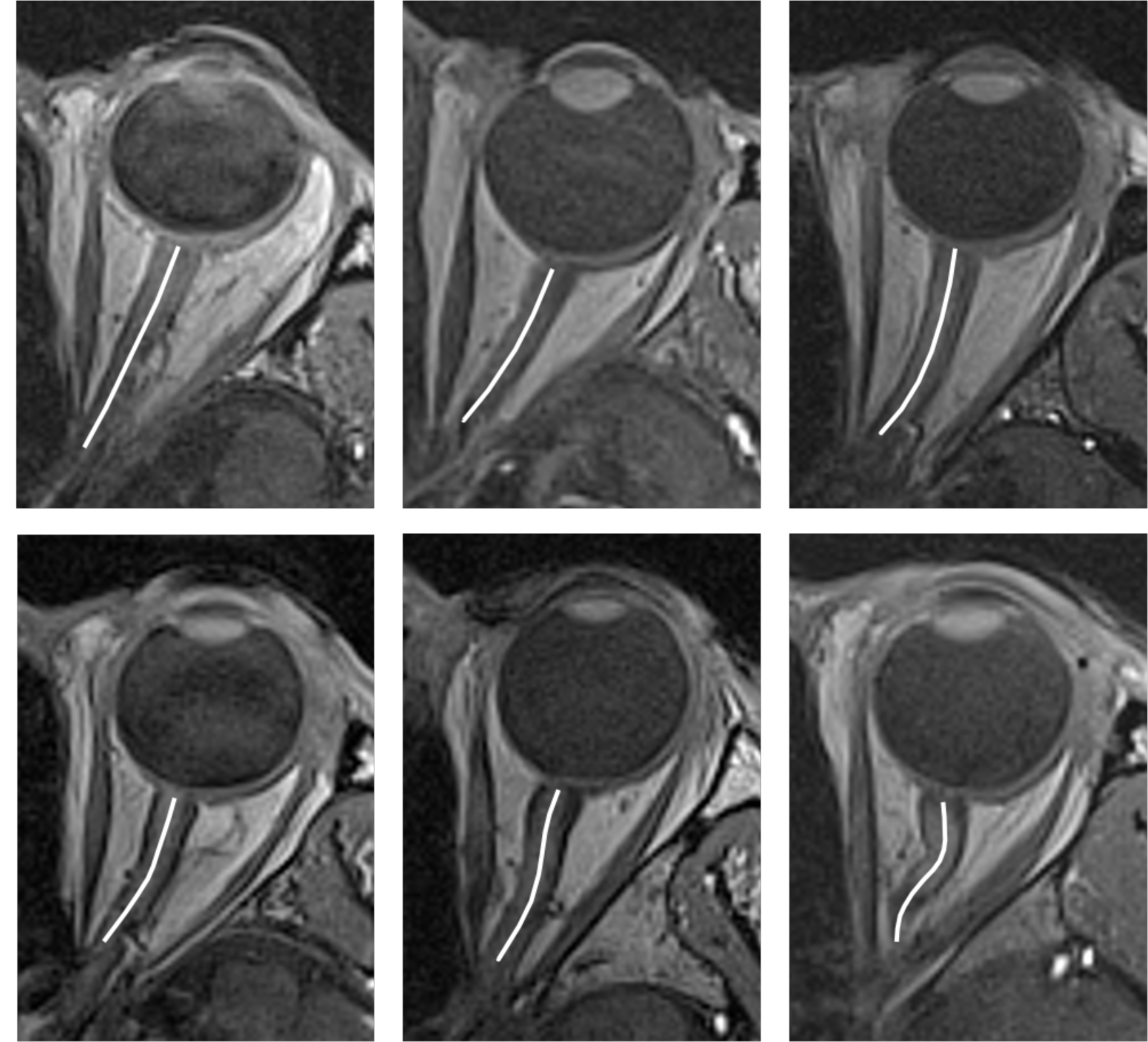
MRI images of the orbital region demonstrate the morphological diversity of the optic nerve (ON). These six figures show examples of ONs displaying varying degrees of curvature, ranging from straight to highly tortuous. The white lines represent the ON middle curve.

The aim of this study was to explore the effects of eye-globe and ON morphologies on gaze-induced ONH deformations, and to examine any potential interactions between morphological parameters and biomechanical parameters of tissues, using parametric FE modeling and design of experiment (DOE).

## METHODS

In this study, we developed a methodology to automatically generate thousands of 3D eye models to study the effects of eye-globe and ON morphologies, as well as tissue biomechanical properties, on gaze-induced ONH deformations. Specifically, a custom-written software (C++) was designed to automatically generate FE models of the eye, each with a set of pre-determined morphological and material parameters. These models were then fed into the FE solver FEBio (Musculoskeletal Research Laboratories, University of Utah, UT, US) to predict gaze-induced ONH deformations. Configurations of all key factors were generated by a DOE approach. The initial DOE analysis evaluated 10 morphological parameters to determine the top 5, which were then combined with the 5 key tissue stiffnesses from previous studies. This resulted in a refined set of 10 parameters, covering both morphology and tissue stiffness, for a follow-up DOE study. Since adduction is known to induce significant ONH deformations compared to abduction,^6,7,9^ an adduction of 13° was chosen for each model, as employed in our previous work. The response of each model was characterized by the magnitude of the effective strain within the LC. Below is a detailed description of the methodology.

### Geometry and Biomechanical Properties of the Baseline FE Model

A whole-eye FE model was established, including the sclera, choroid, prelaminar neural tissue, LC, ON, pia mater, dura mater, orbital fat-muscle complex (OFM) and orbital bone. The baseline geometric parameters of eye global tissues were set to averaged values reported in the literature, shown in **Table 1**. To maintain simplicity, we opted to combine and simulate the extraocular muscles and the orbital fat as a unified entity referred to as OFM. Only half of the eye was reconstructed because the FE model was assumed to be symmetric about a transverse plane passing through the center of the eye globe (**Figure 2**).

**Figure 2.**
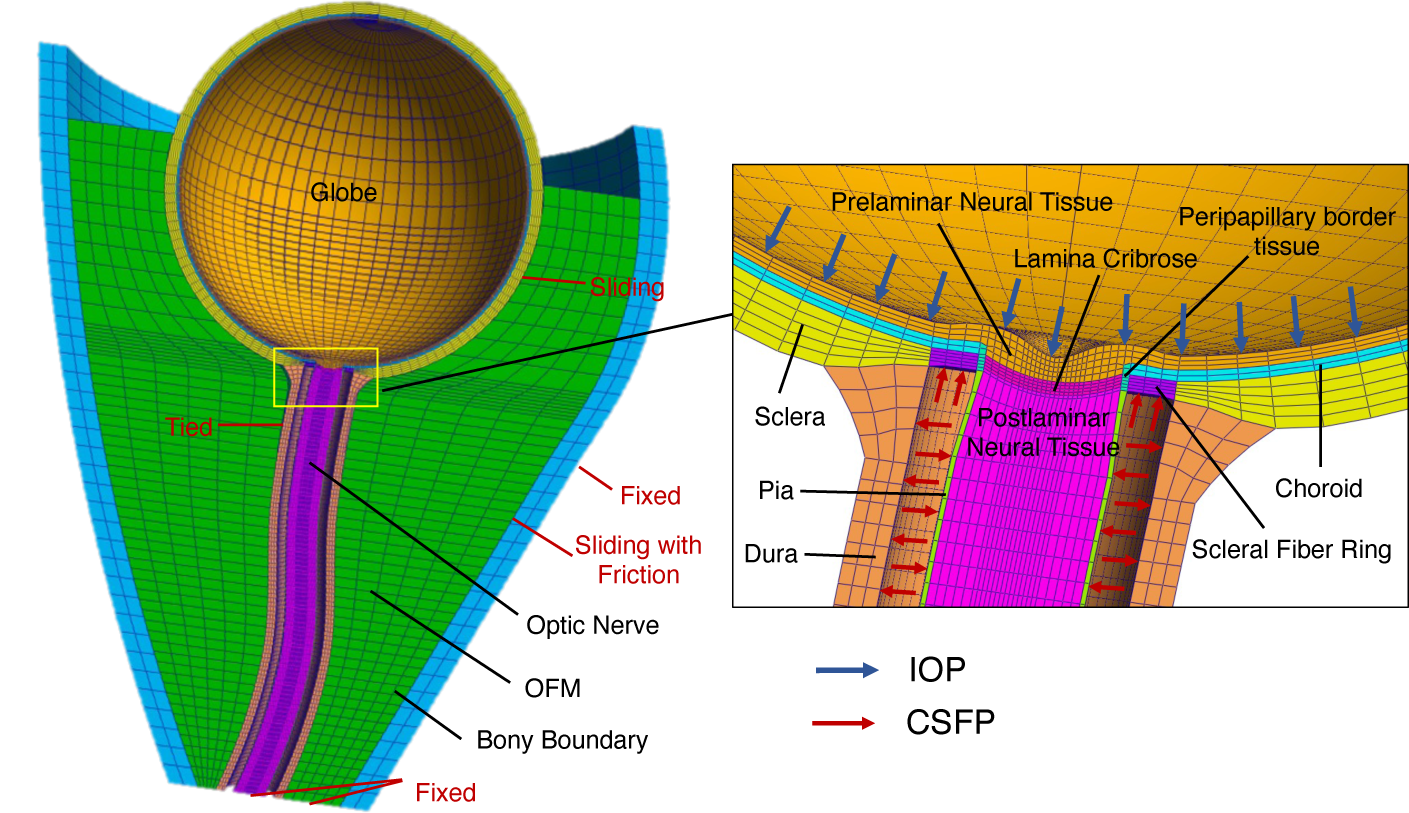
Left panel shows the reconstructed geometry and FE mesh of the eye movement model with boundary conditions and tissue connections. Right panel shows an enlarged view of the detailed ONH region (sclera, scleral fiber ring, the peripapillary border tissue, choroid, Bruch’s membrane, lamina cribrosa, neural tissues, pia and dura) illustrating the IOP and CSFP applied to each model in the primary gaze position.

**Table 1.** Morphological factors and their ranges.

| **Morphological factors** | **Baseline** | **Low** | **High** | **References** |
| --- | --- | --- | --- | --- |
| Dural radius, mm | 2.17 | 1.74 | 2.61 | Vaiman et al.<sup>56</sup> |
| Scleral thickness, mm | 0.996 | 0.7968 | 1.19 | Norman et al.<sup>57</sup> |
| Choroidal thickness, mm | 0.134 | 0.1072 | 0.1608 | Jiang et al.<sup>58</sup> |
| Retinal thickness, mm | 0.249 | 0.1992 | 0.29 | Alamouti<sup>59</sup> |
| ON radius, mm | 1.333 | 1.0664 | 1.5996 | Sigal et al.<sup>60</sup> |
| Pial thickness, mm | 0.06 | 0.048 | 0.072 | Sigal et al.<sup>60</sup> |
| Prelaminar tissue depth, mm | 0.33 | 0.264 | 0.396 | Bowd et al.<sup>61</sup> |
| LC depth, mm | 0.3 | 0.24 | 0.36 | Tun et al.<sup>51</sup> |
| PBT thickness, mm | 0.083 | 0.0664 | 0.0996 | Jonas et al.<sup>62</sup> |
| ON tortuosity | 1.013 | 1.013 | 1.1 | Wang et al.<sup>15</sup> |

The baseline biomechanical properties were the same as those used in our previous studies.^6,7^ Briefly, both the sclera and LC were modeled as soft tissues reinforced with collagen fibers. Those fibers can exhibit stretch-induced stiffening and they are typically distributed within a 2D plane (following a von-Mises probability distribution). The collagen fibers in the peripapillary sclera surrounding the disc were organized into a ring, while those in the peripheral sclera were organized randomly (as specified by the kf parameter in **Table 2**) and parallel to the anterior scleral surface. The collagen fibers in the LC exhibited lower anisotropy than that in the peripapillary sclera and were aligned radially, extending from the central vessel trunk to the LC insertion sites.^20^ All other tissues were considered either hyperelastic or linear elastic as shown in **Table 2**. Among them, peripapillary border tissue (PBT) is the border tissue of the choroid and sclera ^21^. The peripapillary choroid is separated from the prelaminar neural tissue by a collagenous layer, which constitutes the border tissue of the choroid. Likewise, the scleral flange is separated from the LC by the border tissue of the sclera. Since the biomechanical behavior of the PBT has not yet been reported, we assumed that the PBT shared the same biomechanical properties as those of the pia^22^. All soft tissues were assumed to be incompressible. The orbital wall was considered a rigid body.

**Table 2.** Tissue biomechanical properties.

| Tissue | Constitutive Model | Biomechanical Properties | References |
| --- | --- | --- | --- |
| Sclera | Mooney-Rivlin Von Mises Distributed Fibers | $c_1 = 0.805 \text{ MPa}$<br>$c_3 = 0.0127 \text{ MPa}$<br>$c_4 = 1102.25$<br>$k_f = 2$ (scleral ring)<br>$k_f = 0$ (other region of sclera)<br>$\theta_p$: preferred fiber orientations* | Girard et al.<sup>63</sup> |
| PBT | Yeoh model | $c_1 = 0.1707 \text{ MPa}$<br>$c_2 = 4.2109 \text{ MPa}$<br>$c_3 = -4.9742 \text{ MPa}$ | Wang et al.<sup>6</sup> |
| Choroid | Isotropic Elastic | Elastic modulus = 0.6 MPa<br>Poisson's ratio = 0.49 | Friberg et al.<sup>64</sup> |
| Retina | Isotropic Elastic | Elastic modulus = 0.03MPa<br>Poisson's ratio = 0.49 | Miller<sup>65</sup> |
| Lamina Cribrosa | Mooney-Rivlin Von Mises Distributed Fibers | $c_1 = 0.05 \text{ MPa}$<br>$c_3 = 0.0025 \text{ MPa}$<br>$c_4 = 100$<br>$k_f = 1$<br>$\theta_p$: preferred fiber orientation<sup>§</sup> | Zhang et al.<sup>20</sup> |
| Optic nerve | Isotropic Elastic | Elastic modulus = 0.03MPa<br>Poisson's ratio = 0.49 | Miller<sup>65</sup> |
| Pia | Yeoh model | $c_1 = 0.1707 \text{ MPa}$<br>$c_2 = 4.2109 \text{ MPa}$<br>$c_3 = -4.9742 \text{ MPa}$ | Wang et al.<sup>6</sup> |
| Dura | Yeoh model | $c_1 = 0.1707 \text{ MPa}$<br>$c_2 = 4.2109 \text{ MPa}$<br>$c_3 = -4.9742 \text{ MPa}$ | Wang et al.<sup>6</sup> |
| Fat | Isotropic Elastic | Elastic modulus = 0.027MPa<br>Poisson's ratio = 0.49 | Schoemaker et al.<sup>66</sup> |
| Orbit | Isotropic Elastic | Elastic modulus = 300 MPa<br>Poisson's ratio = 0.49 | Schoemaker et al.<sup>66</sup> |

\*Collagen fibers orientations in the scleral fiber ring were aligned circumferentially around the scleral canal. Fibers in other parts of the sclera were organized randomly. Fiber orientations were assigned to these elements using a custom-written MATLAB code.
§ Collagen fibers orientations in the LC were along the radial direction, extending from the central vessel trunk to the scleral canal.

### Parameterization of Morphological and Biomechanical Properties

The morphology of the FE model was parameterized using 10 factors representing the geometry of the eye globe, ONH, and that of the ON. These factors were: dural radius, scleral thickness, choroidal thickness, retinal thickness, ON radius (excluding the pia and dura), pial thickness, the thickness of PBT,^23^ prelaminar tissue depth, central LC depth, and ON tortuosity (**Figure 3**). Specifically, the thicknesses of the eye globe tissues (sclera, choroid, and retina) were modified by adjusting the distance between each tissue’s boundaries and the fixed sclera-choroid interface, while maintaining the thickness of other tissue unchanged. For the ON tissues (pia and PBT), a similar approach was used where the inner surface of each specific tissue was fixed and the outer surface were altered to vary its thickness. The radii of the dura and ON were adjusted by changing their distance from the central axis of the ON and ONH. ON tortuosity was altered by adjusting the positions of three control points along its central path. Refer to **Supplementary Material A-1** for more details on how the morphological parameters of the eye globe and ONH were varied.

**Figure 3.**
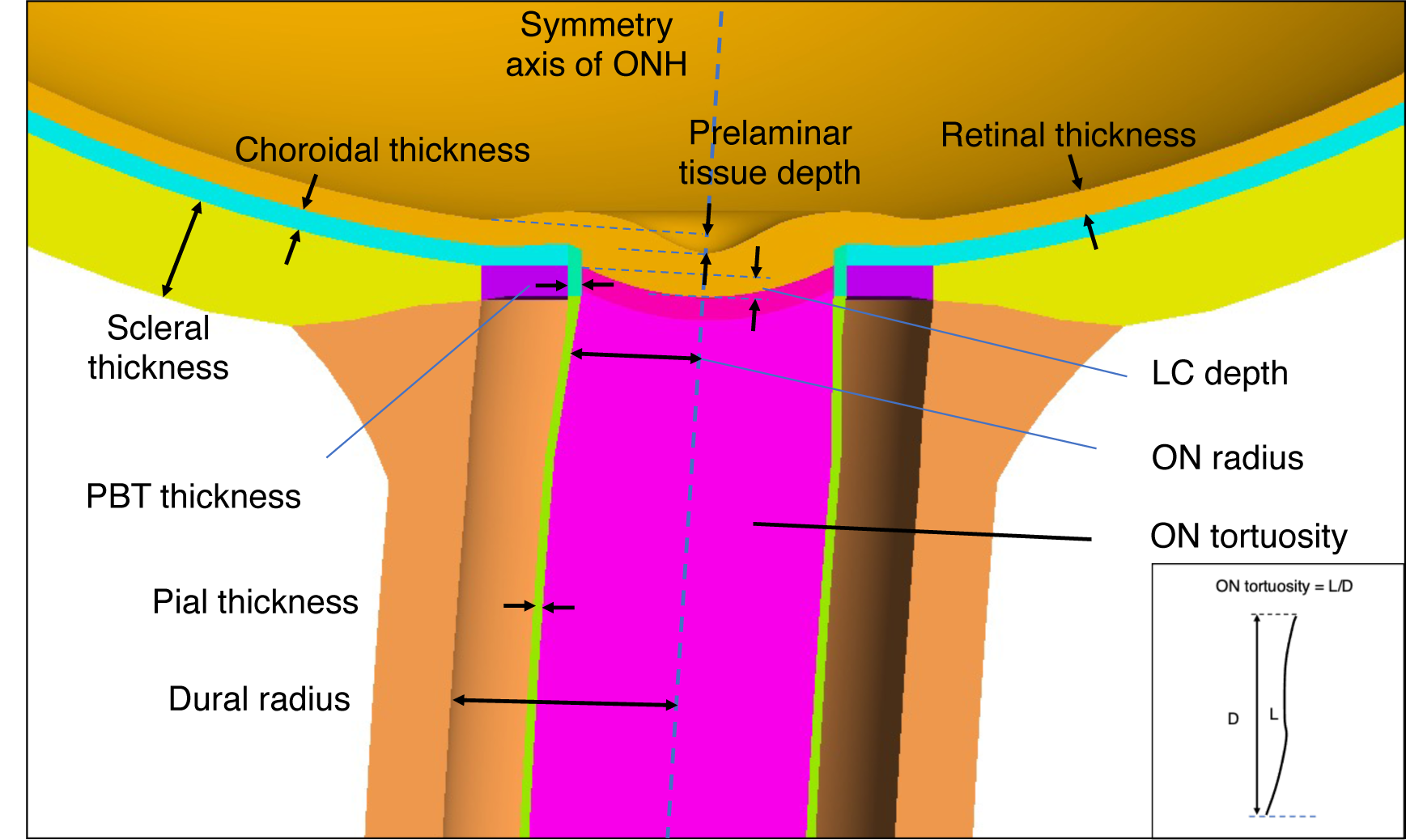
Input factors defining the parametric FE model geometry (only the ONH region of the entire eye is shown). See Table 1 for the ranges of input factors. The blue dashed line represents the symmetry axis of ONH.

The biomechanical properties of each tissue were directly modified in the input file of the FE model.

### Contact Definitions, Boundary and Loading Conditions

Contacts between tissues were the same as those defined in our previous study.^6,7^ Briefly, the OFM and the dura were tied together; the OFM was able to slide over the bony orbital margin with a friction factor of 0.5; the cornea-scleral shell and the OFM had a sliding contact with no friction to mimic the Tenon’s capsule enveloping the eyeball. Rigid contact was assigned between the horizontal rectus muscle insertions and a ‘non-physiological’ rigid body. This latter had a center of mass at the center of the eyeball, which was constrained with a prescribed rotation to simulate an adduction of 13°. For boundary conditions, the OFM and ON were fixed at the orbital apex to mimic the fibrous adhesion of those tissues to the optic canal. The orbital bone was also held in place by fixing its outer margin. In addition to an adduction of 13°, an IOP of 15 mm Hg was applied to the surface of the retinal and prelaminar tissues and a CSFP of 12.9 mm Hg was applied within the subarachnoid space of the ON. Loading was applied in two steps: first, IOP and CSFP were applied and maintained, then an adduction of 13° was applied. All contact patterns, boundaries and loading conditions are illustrated in **Figure 2**.

### FE Simulations, Post-processing and Output Measurement

All FE models were consistently meshed with 73,922 nodes and 62,821 elements, including 62,521 8-node hexahedra and 300 6-node pentahedra elements. All tissues were bonded by shared nodes at tissue boundaries (**Figure 2**). The mesh density was numerically validated through a convergence test as described in our previous study.^6^ A single model required about 20 minutes to solve on a desktop workstation (Intel Xeon Silver 4114 CPU @ 2.20GHz, 32GB of memory).

The preprocessing Matlab script executed FEBio to solve the generated models and outputted the Lagrange strain tensors for each step and the volumes of all LC elements into a text file. Effective strains were calculated from the principal components of the Green Lagrange strain tensor. To isolate the specific effect of eye movement, effective strains of each LC element after the first load step (with only IOP and CSFP) and those after the second load step (following eye movement) were extracted. The differential, termed “delta effective strain”, was then calculated for each element. This delta effective strain for each LC element was multiplied by its volume, and these values were summed and divided by the total LC volume, yielding the volume-weighted mean LC delta effective strain. For simplicity, this will be referred to as the LC effective strain in the subsequent manuscript.

### DOE of Morphological Factors

For the 10 morphological parameters (**Figure 3**), a two-level full factorial design was used, resulting in a total of 1,024 models. This comprehensive design allowed us to assess the effects of the main factors and any potential interactions among them. The low and high levels of all morphological parameters (excluding ON tortuosity) were set by varying them by 20% around their baseline values (**Table 1**). For simplicity, we varied ON tortuosity within a narrower range, from 1.013 (low level) to 1.1 (high level), as these values have typically been observed in MRI studies. Detailed information on the morphological parameters for each model is available in the **Supplementary Material B-Sheet1.**

### DOE of Both Morphological and Biomechanical Factors

We conducted another DOE analysis to examine the potential interactions between morphological parameters and tissue stiffnesses. This analysis included five key morphological parameters identified from the initial DOE analysis and the stiffnesses of five tissues (LC, ON, sclera, pia and dura) informed by our previous study.^6,7^

In this DOE analysis, high and low levels of the biomechanical properties of LC, ON, sclera, pia and dura were set by varying the material constants by 20% around their baseline values (see **Table 3** for the exact values). The variation in morphological parameters was consistent with the initial DOE analysis. A two-level full factorial design was employed, resulting in 1,024 models. Detailed information on the morphological parameters and tissue stiffnesses for each model is available in the **Supplementary Material B-Sheet2.**

**Table 3.**
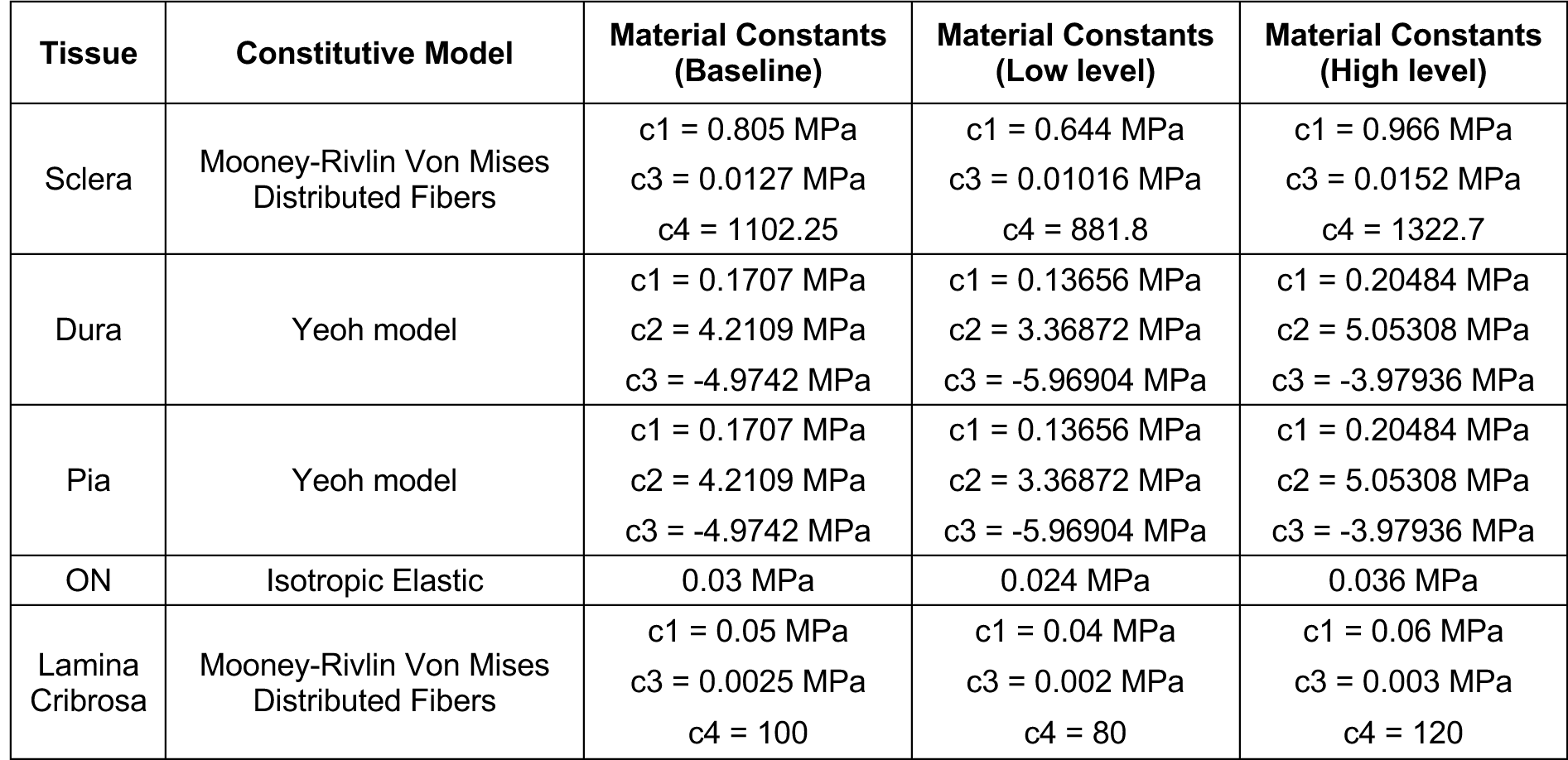
Biomechanical properties of five tissues and their ranges.

### Statistical Analysis

All statistical analyses were carried out in Minitab (release 20, Minitab, LLC, Pennsylvania, USA). A P-value of less than 0.05 was considered statistically significant.

For each model, we reported gaze-induced LC effective strain as responses. Main effects and interaction effects were reported, and the significance of factors was tested and ranked as detailed below. The main effect indicates the average change in the response when a factor’s level shifts from low (512 models) to high (512 models), in which other factors vary between both levels. Interaction effects refer to how the influence of one factor on the response changes depending on the level of another factor. Essentially, it examines whether the combined impact of two factors differs from the sum of their individual effects.

Analysis of Variance (ANOVA) was used to assess the significance of individual factors or interactions when they vary from low to high levels. It is important to note that in DOE, the use of ANOVA is not limited by the number of levels a factor has. Even when a factor is set at just two levels, as in this study, ANOVA remains well-suited for assessing the factor’s significance on the response variable and continues to be the standard method in Minitab. This is because ANOVA focuses on partitioning the total variability in the data into components attributable to different sources, including the main effects of factors and their interactions. As ANOVA was performed individually for all factors and their possible interactions, we applied the Bonferroni correction to our p-values to account for the increased risk of Type I errors due to multiple comparisons.

In the DOE analysis, a linear relationship between the factors (and their interactions, if included) and the response is assumed and liner models were fitted with these factors as independent variables. In this context, the R^2^ value indicates how well the linear model fits the experimental data, essentially assessing how effectively the factors explain the variation in the response. Statistically significant factors were further ranked based on the magnitude of change they induced in the response variable (equals to the absolute value of the regression coefficients), facilitating the identification of the most influential factors and interactions.

## RESULTS

### The Effects of 10 Morphological Parameters

The average LC effective strain (i.e., delta strain after removing the effects of IOP and CSFP) across all models was 0.031. In the DOE analysis, we examined 55 factors, comprising 10 main factors and 45 pairwise interactions. Out of these, 25 factors were statistically significant (p < 0.05). Among the significant factors, only five factors contribute to more than 1% of the total effects: ON tortuosity, dural radius, ON radius, scleral thickness and LC depth. A linear regression model showed that these five factors accounted for 96.69% of the total effects in the responses. Details on the statistically significant factors with less than 1% impact on the total effects are available in **Supplementary Material B-Sheet3**.

Larger ON tortuosity, scleral thickness, ON radius and LC depth decreased LC strains following eye movements, while a lager dural radius increased LC strains. **Figure 4** illustrates the magnitude and trend of the effects. **Figure 5** shows the morphology of the undeformed and deformed FE models and color-coded strains (ONH and LC) in models with low and high levels of these five factors. In the baseline model, the mean LC effective strain caused by 13° adduction was 0.042. The mean LC effective strains for models with larger ON tortuosity, dural radius, ON radius, scleral thickness and LC depth were 0.026, 0.054, 0.026, 0.035, and 0.037, respectively.

**Figure 4.**
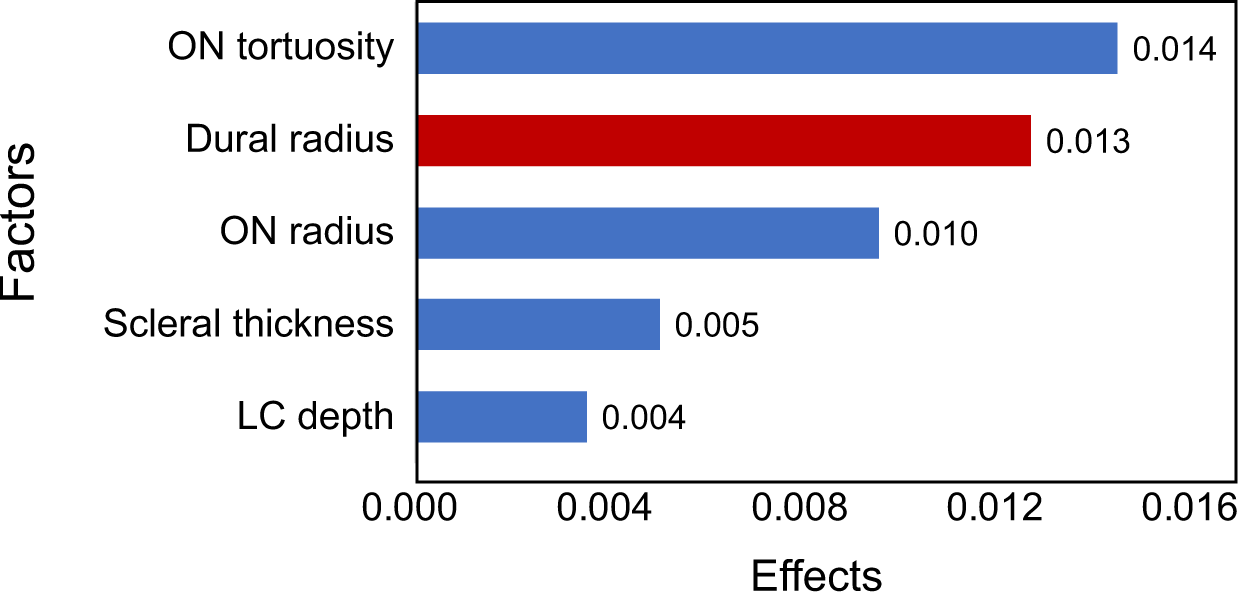
Ranking of the effects of morphological factors (only the five most significant factors contributing to more than 1% of the total effects were shown) on the mean effective strain of LC. A longer bar indicates a more significant effect when varying parameters from a low to a high level. Blue bars indicate positive effects (strain reduction) and red bars indicate negative effects (strain increase).

**Figure 5.**
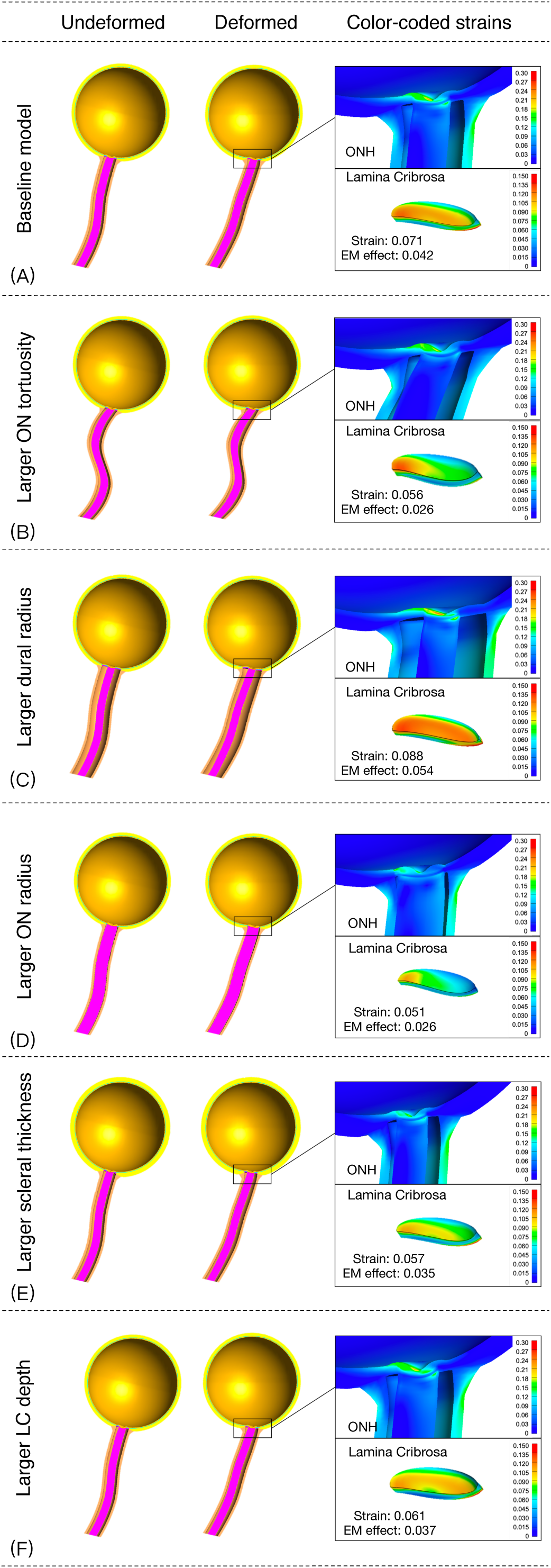
ONH deformations induced by an adduction of 13° with the five main factors (ON tortuosity, dural radius, ON radius, scleral thickness and LC depth) at their low and high levels, respectively. The enlarged views of the ONH and LC show the color-coded effective strain. In the enlarged views, “Strain” represents the total LC effective strain induced by IOP, CSFP and eye movement, and “EM effect” represents the mean LC delta effective strains after removing the effects of IOP and CSFP. Note that the LC deformations were exaggerated 5 times for illustration purposes.

### The Effects of 5 Morphological Parameters and 5 Tissue Stiffnesses

A total of 55 factors, including main factors and their interactions, were examined; 39 of these were statistically significant (p<0.05). The factors that individually contributed to more than 1% of the total effects, in descending order, are ON tortuosity, dural radius, ON radius, dural stiffness, scleral thickness, LC stiffness, LC depth, the interaction between the dural radius and the dural stiffness, together accounting for 95.52% of the total effects in the responses. For detailed information on factors that were statistically significant but had an effect of less than 1% of the total effects, refer to **Supplementary Material B-Sheet4**.

Figure 6 illustrates the magnitude and trend of the effects. The trend for the 5 morphological parameters are consistent with the results from the above morphological DOE analysis. For biomechanical properties, a stiffer dura increased LC strains, whereas a stiffer LC reduced strains. The most pronounced interaction occurred between dural radius and dural stiffness. Specifically, an increase in LC strain associated with a larger dural radius was found to be amplified when combined with a stiffer dura.

**Figure 6.**
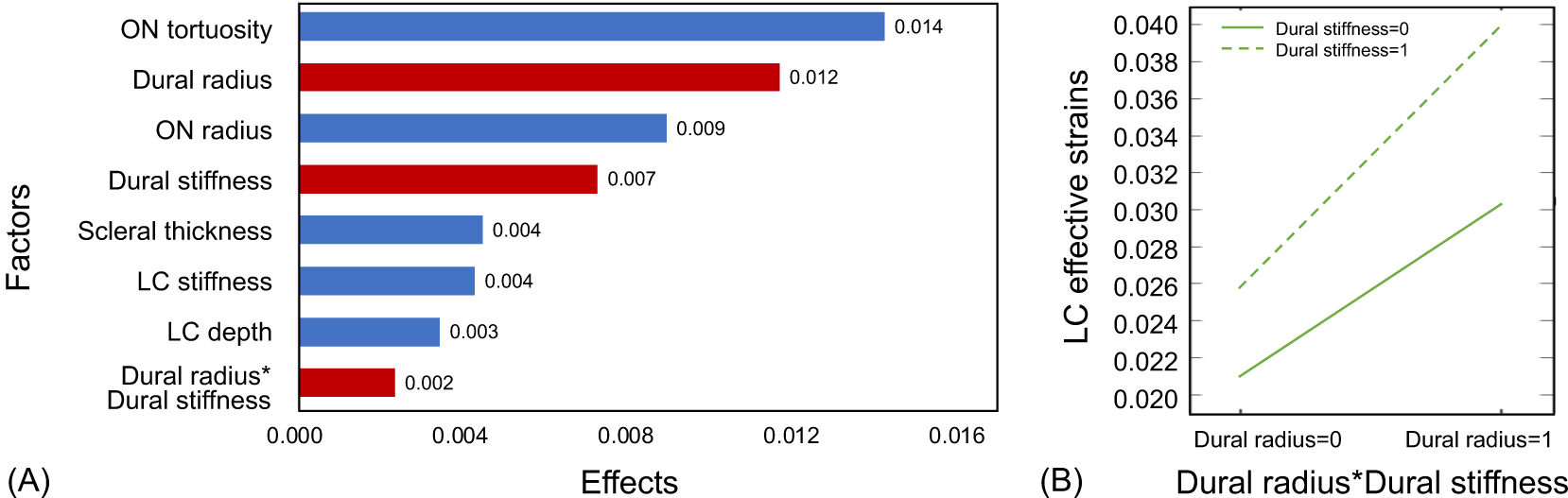
(A) Ranking of the effects of morphological factors, tissue stiffnesses and their interactions (factors contributing to more than 1% of the total effects were shown) on the mean effective strain of LC. A longer bar indicates a more significant effect when varying parameter from a low to a high level. Blue bars indicate positive effects (strain reduction) and red bars indicate negative effects (strain increase). (B) The interactions between dural radius and dural stiffness. When dural stiffness is at the low level, LC effective strain increased by 0.0093 (from 0.0210 to 0.0303) with an increase in dural radius. At the high level of dural stiffness, LC effective strain increased by 0.0140 (from 0.0259 to 0.0399) with the increase of dural radius. 0, low level; 1, high level.

## DISCUSSION

In this study, we developed a parametric FE model and studied the effects of eye-globe and ON morphologies, as well as tissue stiffnesses on ONH deformations during eye movements. Our models demonstrated that ON tortuosity, dural radius, ON radius, scleral thickness and LC depth were the five main morphological factors that significantly affect gaze-induced ONH deformations. These parameters retained their significance in a combined analysis with tissue stiffnesses. We also observed a significant interaction between the dural radius and dural stiffness, proving to be a considerable factor in the ONH’s response to eye movement.

### A Larger Dural Radius Increased LC Strains During Eye Movements

Our study found that a larger dural radius (i.e., the optic nerve sheath inner radius) leads to higher LC strains, similar to the effect of a stiffer dura. During eye movements, peripapillary tissues are sheared in the transverse plane by the optic nerve sheath, resulting in significant deformations.^7^ An increased dural radius would tend to restrict eye movements by exerting a larger pulling force onto the ONH, as evidenced by the calculated traction force from the FE models (**Supplementary Material A-2**). The potential for such forces to cause axonal death in glaucoma requires further investigation.

In this study, it is important to acknowledge that variations in the dural radius were accompanied by changes in the length of the scleral flange, as the insertion point of the dura into the sclera was altered. Consequently, an increase in dural radius resulted in an enlargement of the scleral flange in the model. Although this relationship aligns with anatomical observations, where the dural radius and scleral flange size are positively correlated,^4,24^ this confounding factor complicates the interpretation of the effect of a larger dura radius. Specifically, a larger scleral flange could potentially indicate a weaker ONH, which is more susceptible to deformation. To dissect the potential confounding impacts of an increased scleral flange and dural radius, we conducted additional simulations (**Supplementary Material A-3**). In these models, we modified the dural and ON radius without varying the scleral flange size. Keeping the scleral flange size constant, we observed that an increase in dural radius (along with a concurrent increase in ON radius, which can actually reduce LC strain) led to an increase in LC strains. This approach allowed us to confirm that an increased dural radius indeed contributes to higher LC strains. Computed tomography (CT) studies have shown that the optic nerve sheath diameter (ONSD) is significantly larger in patients with normal tension glaucoma (NTG) compared to healthy controls^25,26^, suggesting that NTG eyes may exhibited more ONH deformation due to eye movement. However, other studies have found no significant difference in ONSD between NTG and healthy controls,^27^ and that NTG subjects may have a smaller ONSD owing to a lower CSFP.^28,29^ These conflicting findings could be due to ethnic differences or underlying differences in the pathogenesis of various NTG subtypes. Our previous studies demonstrated a negative correlation between IOP-induced ONH strain and retinal sensitivity in high tension glaucoma subjects^17^, whereas NTG subjects showed a stronger correlation with gaze-induced ONH deformation^30^. Given these results, it is quite possible that NTG itself may have different subtypes, with some subjects being sensitive to IOP and others sensitive to gaze. The role of morphological differences in the dura in these variations remains unexplored, and further studies are warranted.

### A Large ON Tortuosity Decreased LC Strains During Eye Movements

Our study suggested that increased ON tortuosity may lead to lower LC strains during eye movements. A relatively taut ON has the potential to exhibit rapid straightening during eye movements and thus exert more force on the ONH tissue, a phenomenon observed in our FE models (**Supplementary Material A-2**). In a previous study involving a small Chinese population, we found that ONs in glaucoma subjects (mean IOP: 26.4 ± 4.6 mmHg) were tauter than in normal controls (mean IOP: 15.3 ± 3.6 mmHg). This smaller ON tortuosity in glaucoma subjects may exert more force on the ONH tissues during eye movements, indicating a potential risk factor for glaucoma.^15^ However, another study^31^ reported that ON path redundancy was greater in NTG than in normal controls in primary gaze and abduction. The discrepancy between those two studies may be attributed to: 1) differences in subjects (high-tension glaucoma subjects of Chinese ethnicity versus NTG subjects of unknown ethnicity); 2) small sample sizes; or simply; 3) differences in methods to assess ON tortuosity.

A recent study reported that ON tortuosity in highly myopic subjects was significantly larger than that in emmetropic controls.^16^ It would seem that, in high myopia, the ‘slack’ ON (i.e., increased ON tortuosity) might act as a protective mechanism against ONH deformations. However, it is crucial to acknowledge that high myopic subjects with more tortuous ONs might still be susceptible to greater ONH deformations during eye movements due to other influencing factors. For instance, in high myopic eyes, the ON-globe junction must travel a longer distance for the same amount of eye movement compared to normal eyes. This is due to the extreme elongation of high myopic eyes, which can exhaust the redundancy in ON tortuosity. Additionally, the weakened structural stiffness of the sclera and other ONH structures in high myopia could make them more susceptible to ONH deformations.

### A Thicker Sclera Decreased LC Strains During Eye Movements

Our study showed that scleral thickness significantly affects LC strains, with a thicker sclera associated with lower LC strains during eye movements. This finding aligns with other studies examining the effects of factors on IOP-^32,33^ and CSFP-^5^ induced LC strains. Since other ONH tissues are relatively compliant compared to the sclera, scleral deformation induced by eye movement can be directly transmitted to surrounding tissues, suggesting that eyes with a thinner sclera may be more sensitive to eye movements. In case of high myopia, scleral thickness decreases significantly with increasing axial length.^34^ In severe cases, scleral thickness can be as low as 31% of that in normal subjects^35^, potentially leading to the development of staphylomas. The reduced scleral thickness in high myopia could result in large LC deformations and increasing susceptibility to ONH damage during eye movements. However, as discussed above, ONH deformations in high myopic eyes are also affected by axial length and ON tortuosity. Further studies are warranted to investigate the interactions of these factors in high myopia.

### Other Factors Affecting LC Strains During Eye Movements

Our study showed that a larger ON radius (i.e., the ON parenchyma, excluding the dural and pial sheaths) had a protective effect, resulting in smaller LC strains. There is a typical shear deformation due to ON traction, with clear temporal pulling from the dura in adduction. We speculated that a larger ON radius might possibly provide more mechanical support to the LC during ON traction, which could potentially lead to smaller LC strains. ON radius is associated with disc size. Previous studies have reported conflicting results regarding the relationship between disc size and glaucoma. Some studies^2,36,37^ demonstrated that a larger optic disc size is associated with higher glaucoma susceptibility, while other studies suggested that smaller discs with less space for nerve fibers to travel through increase glaucoma susceptibility^38,39^. There are also studies^40,41^ found no significant correlation between the degree of ON atrophy in glaucoma and disc size. Note that disc size measured in these studies were not equal to the ON size posterior to LC. Direct measurement of ON size with MRI imaging^42,43^ have shown that the ON radius of glaucoma subjects was smaller than that of normal controls, which was significantly correlated with retinal nerve fiber layer thickness thinning and perimetric loss. A recent study suggested that myopes also tended to have smaller ONs.^44^ It is worth noting that histologic studies showed that optic atrophy led to a smaller retrobulbar ON^45^, suggesting that ON diameter may correlate with the extent of optic atrophy and a smaller ON diameter in glaucoma subjects might be the consequence of retinal ganglion cells (RGC) apoptosis^31^. The link between ON size and glaucoma needs further exploration.

Our study also revealed that a larger LC depth resulted in smaller gaze-induced LC strains. In this study, a greater LC depth corresponds to a more curved LC. Previous studies^46–49^ showed that the LC depth and LC curvature were significantly larger in POAG eyes than in healthy eyes. These differences could be the consequences of glaucoma.^50^ However, it remains unclear whether an initial larger LC depth/curvature is protective or detrimental in the development of glaucoma. Since LC morphology varies with race, sex, age and axial length^51–54^, the relationship between the LC morphology and gaze-induced LC deformations needs to be further studied.

Lastly, our study reconfirmed the significance of tissue stiffness on gaze-induced LC deformations. A stiffer dura increases LC strains, which is consistent with our previous study.^6^ In addition, LC stiffness also had a strong influence on gazed-induced LC strains, where a stiffer LC resulted in a reduction of LC strains. This is straightforward as a stiffer material will deform less under the same loading condition. This observation is consistent with other studies investigating LC strains induced by IOP^32,33^ and CSFP^55^. However, the effect of tissue stiffness on gazed-induced LC strains is smaller than that of the three main morphologies (ON tortuosity, dural radius, and ON radius; Figure 6 and **Supplementary Material B-Sheet4**).

### The Interactions Affecting LC Strains During Eye Movements

This study revealed significant interactions among various factors, with the most notable being between dural radius and dural stiffness. A larger dural radius tends to increase LC strains, an effect that is amplified with stiffer dura and diminished when the dura is more compliant. These findings underscore the importance of considering individual-specific characteristics such as eye globe and ON morphologies, as well as their biomechanical properties, in assessing the susceptibility of LC deformation during eye movements. Given the complex and multifaceted nature of morphological and biomechanical properties of the ONH, our parametric FE models provided an ideal platform for studying and quantifying the main factors and their interactions in a systematic manner to inform future experimental study design and analysis.

### Limitations

In this study, several limitations warrant further discussion. First, our models only predicted acute ONH deformations during eye movements and could not account for the long-term growth & remodeling processes that are known to take place in ocular tissues.

Second, there are some inherent limitations in a two-level full factorial design. As each factor has only two levels, it cannot account for the nonlinear effect between the factor and the response. Additionally, in a full factorial design, all possible combinations of the factors are tested, resulting in a high number of experiments. This would increase the time and cost to conduct the study. Considering more advanced experimental designs that account for nonlinear effects and minimize the number of experiments can enhance the efficiency and precision of future studies in this field.

Third, the morphological factors in our study included dural radius, but not dural thickness. Previous computational studies and this study have shown that a stiffer dura could significantly increase LC strains during eye movements.^6^ To enhance our understandings, future studies should examine the effect of an increased dural thickness on LC deformation during eye movements.

Fourth, to rank the effects of all morphological factors, we varied these parameters by 20% from their baseline values, as proper physiologic ranges for each parameter are not known. As a result, morphological size variations from larger tissues were higher. A more precise understanding of the physiological ranges for these parameters would be valuable for more accurate assessments in future studies.

Finally, the simplified morphological properties of our models provided a reasonable approximation, allowing us to improve our understanding of ONH biomechanics during eye movements. It will be necessary to update this work as more biomechanical information on eye and orbital tissues becomes available.

### Conclusion

Our parametric finite element models demonstrated that ON tortuosity, dural radius, ON radius, scleral thickness and LC depth were the five most important morphological factors influencing gaze-induced ONH deformations. Additionally, the stiffnesses of dura and LC were the most important biomechanical factors influencing gaze-induced ONH deformations, and the interactions between dural radius and the dural stiffness was significant. Our study provides an ideal platform for studying and quantifying the main factors and interactions between factors to inform experimental design and analysis. Further experimental and clinical studies are needed to explore the role of effect of individual-specific characteristics on gaze-induced ONH deformations in ocular diseases, such as myopia and glaucoma.

## ACKNOWLEDGMENTS

Acknowledgement is made to (1) the National Natural Science Foundation of China (12272030, 12002025), (2) the donors of the National Glaucoma Research, a program of the BrightFocus Foundation, for support of this research (G2021010S [MG]), (3) NMRC-LCG grant ‘TAckling & Reducing Glaucoma Blindness with Emerging Technologies (TARGET)’, award ID: MOH-OFLCG21jun-0003 [MG], (4) the “Retinal Analytics through Machine learning aiding Physics (RAMP)” project that is supported by the National Research Foundation, Prime Minister’s Office, Singapore under its Intra-Create Thematic Grant “Intersection Of Engineering And Health” - NRF2019-THE002-0006 awarded to the Singapore MIT Alliance for Research and Technology (SMART) Centre [MG].

## Notes

### Competing Interest Statement

The authors have declared no competing interest.

### Summary of Updates

Figure 5 revised；acknowledgments updated.

