## Supplementary Material A for "Effect of eye globe and optic nerve morphologies on gaze-induced optic nerve head deformations"

### Supplementary Material A-1

The thicknesses of the eye globe tissues (sclera, choroid, and retina) were altered by adjusting the distance between each tissue's boundaries and the fixed sclera-choroid interface (red dashed line in **Figure S1**), while maintaining the thickness of other tissue unchanged. For the optic nerve tissues, the thicknesses of pia and PBT were changed in a similar approach, where the inner surface of each specific tissue was fixed and the outer surface were altered to vary its thickness. The radius of the dura and ON were adjusted by changing their distance from the central axis of the ON and ONH. **Figure S1** illustrates the reference boundaries (the starting points of the black arrows) and the increase of tissue thickness/size following the directions indicated by the black arrows.

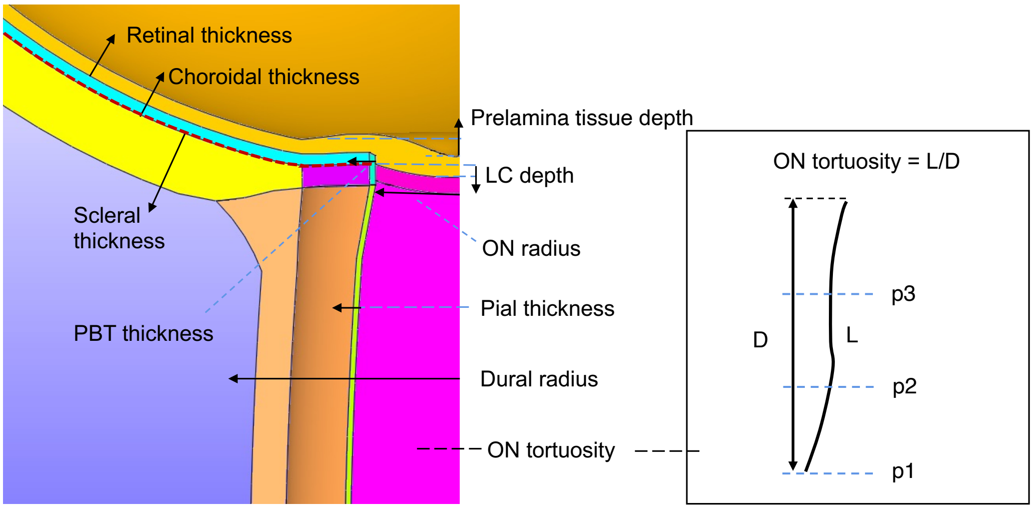


**Figure S1.** Illustration of morphological parameter variations in the finite element model. The red dashed line indicates the fixed sclera-choroid interface. The reference boundaries are indicated by the starting points of the black arrows. Tissue thickness/size was varied by altering the boundary opposite the reference boundary, with increases indicated by the arrow direction and decreases in the opposite direction.

The morphology of the ON was determined by three control points (p1, p2, p3 in **Figure S1** and **S2**), representing the end of the ON at the orbital apex, 30% along the ON and 65% along the ON. The coordinates of these three points are controlled by the deviation distances of the ON (depicted as curve **L**, a black line in **Figure S2**) with respect to a straight ON (depicited as curve **D**, a red line in **Figure S2**) in the naso-temporal direction. ON tortuosity was altered by adjusting the positions of three control points along its central path. The high and low levels of ON tortuosity were set as 1.1 and 1.013, in which the deviation distances of these three points were (0mm, 2.8mm, -2.2mm) and (-1.4mm, 0.5mm, -0.4mm), respectively.

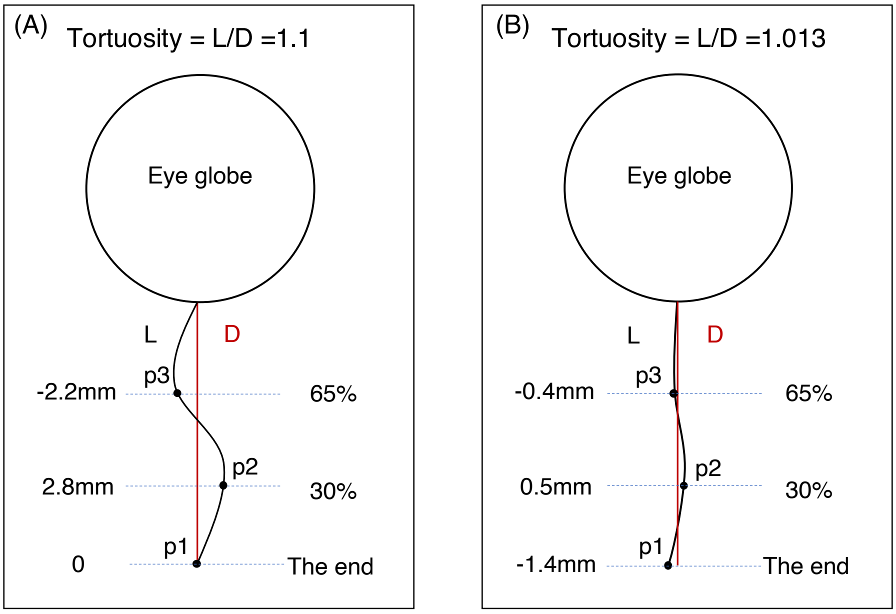


**Figure S2.** ON tortuosity at the high and low levels. (A) The high level of ON tortuosity were set as 1.1, in which the deviation distance of three point were 0,

2.8mm, -2.2mm, respectively. (B) The low level of ON tortuosity were set as 1.013, in which the deviation distance of three point were -1.4mm, 0.5mm, -0.4mm, respectively.

### Supplementary Material A-2

Traction forces were measured along the deformed optic nerve direction defined from the ONH center to the orbital apex as illustrated by the black arrows in **Figure S3**. Specifically, for each dural element adjacent to the sclera, the normal stress along the deformed optic nerve direction was derived from the stress tensor of that element. This normal stress was then multiplied by the area of the middle cross-section of the element to obtain the force value. The forces of all these elements were summed and reported as the traction force. This calculation method was the same as in our previous study[1].

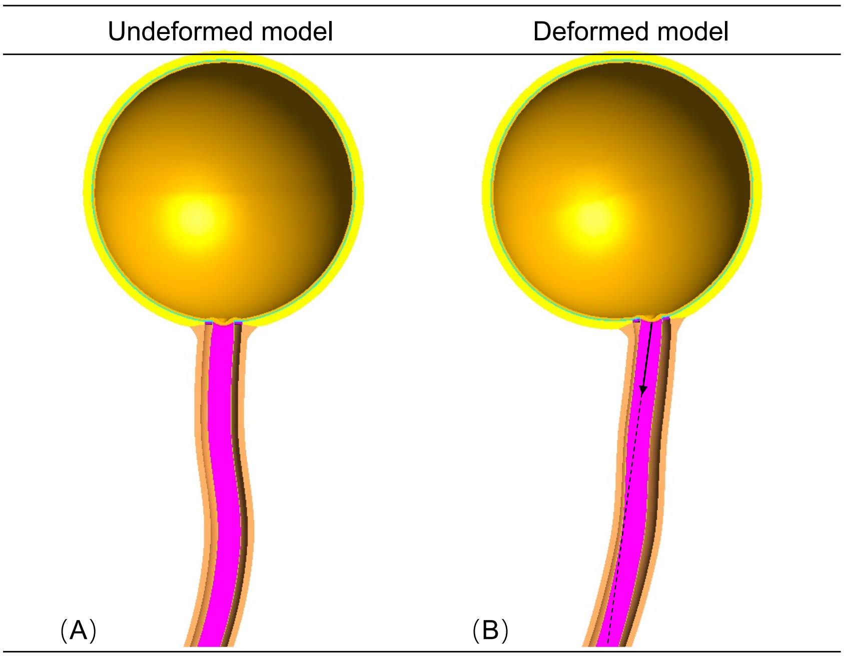


**Figure S3.** Schematic diagram of the undeformed model (A) and deformed model (B). Black arrows superimposed on the optic nerve indicate the directions of the reported traction forces during eye movements.

In the Design of Experiment (DOE) for morphological factors, the three most significant factors affecting ON traction force were ON tortuosity, dural radius, and scleral thickness ($p<0.05$). **Figure S4** illustrates the magnitude and trend of these effects. Larger ON tortuosity and scleral thickness reduced traction force, while a larger dural radius increased the traction force. An increased dural radius would tend to restrict eye movements by exerting a larger pulling force onto the ONH. A relatively taut ON has the potential to exhibit rapid straightening during eye movements and thus exert more force on the ONH tissue.

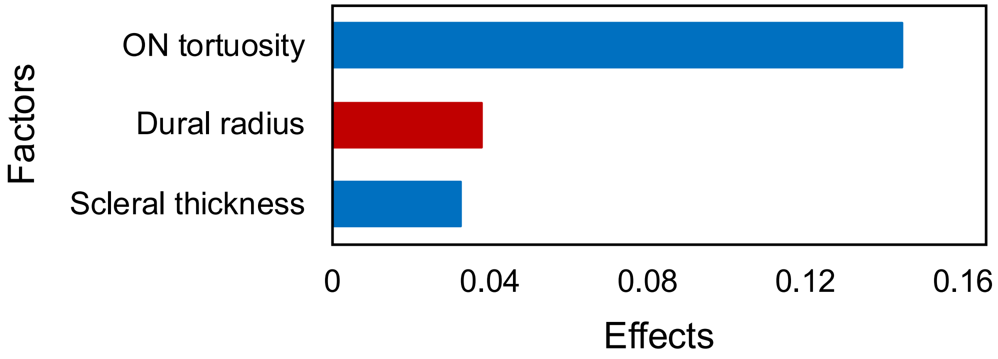


**Figure S4.** Ranking of the three most significant factors (ON tortuosity, dural radius and scleral thickness) affecting ON traction force. The horizontal bars represent the effects of factors. A longer bar indicates a more significant effect when varying parameter from a low to a high value. Blue bars indicate positive effects (ON traction force reduction) and red bars indicate negative effects (ON traction force increase).

### Supplementary Material A-3

To dissect the potential confounding impacts of an increased scleral flange and dural radius, we conducted additional simulations. In these simulations, we increased the dural radius while simultaneously increasing the ON radius, all the while keeping the scleral flange size constant (refer to **Figure S5**).

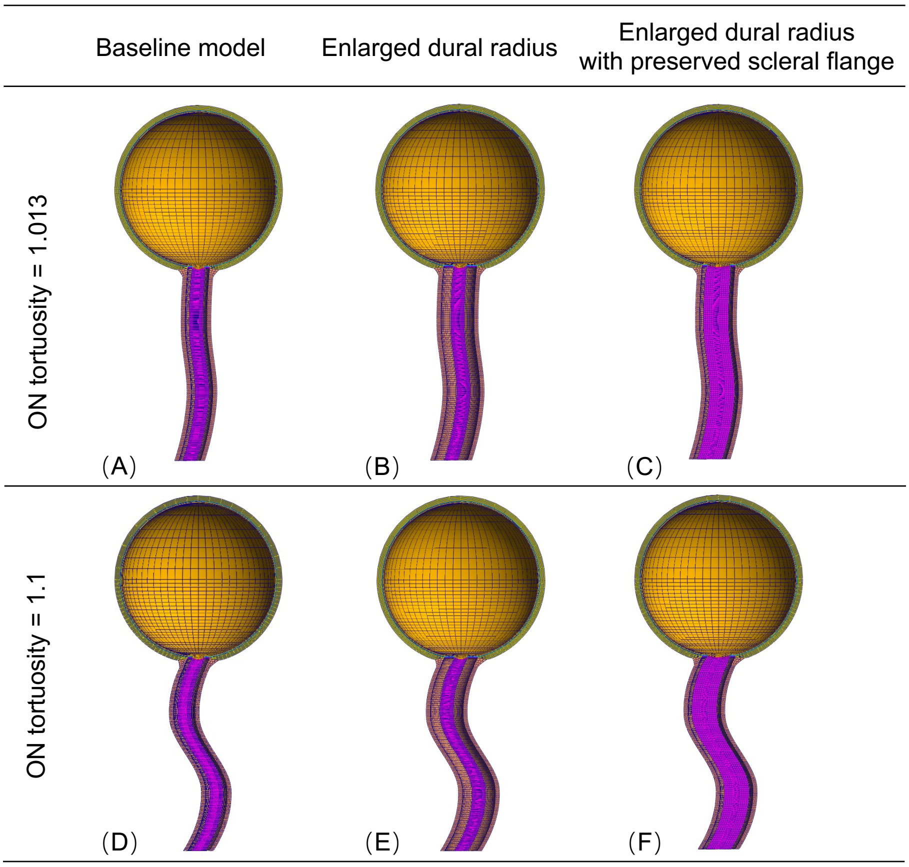


**Figure S5.** Upper row: At low ON tortuosity (1.013), showing the undeformed baseline model (A), the model with a larger dural radius (B), and the model with both a larger dural radius and preserved scleral flange size (C). Lower row: At high ON tortuosity (1.1), showing the undeformed baseline model (D), the model with a larger dural radius (E), and the model with both larger dural radius and preserved scleral flange size (F).

Keeping scleral flange size unchanged, the increase of dural radius (along with a concurrent increase in ON radius) led to the increase of gaze-induced LC strains (**Table S1**). Notably, an increase in ON radius had the effect of reducing LC strains. Consequently, this approach provided confirmation that an increased dural radius indeed contribute to higher LC strains.

**Table S1.** LC strains of models with different dural radius, scleral flange, ON radius and ON tortuosity (ON T).

| **Models** | **Dural radius (mm)** | **ON radius (mm)** | **Scleral flange (mm)** | **LC strains (ON T=1.013)** | **LC strains (ON T=1.1)** |
|---|---|---|---|---|---|
| **Baseline model** | 1.740 | 1.066 | 0.626 | 0.037 | 0.023 |
| **Enlarged dural radius** | 2.610 | 1.066 | 1.496 | 0.046 | 0.031 |
| **Enlarged dural radius with preserved scleral flange** | 2.610 | 1.936 | 0.626 | 0.039 | 0.025 |
